# *PopGenPlayground:* a population genomics analysis pipeline

**DOI:** 10.1101/2024.02.27.582400

**Authors:** Walter W. Wolfsberger, Khrystyna Shchubelka, Olga T. Oleksyk, Yaroslava Hasynets, Silvia Patskun, Mykhailo Vakerych, Roman Kish, Violeta Mirutenko, Vladislav Mirutenko, Coralia Adina Cotoraci, Calin Pop, Olimpia Neagu, Cornel Baltă, Hildegard Herman, Paula Mare, Simona Dumitra, Horatiu Papiu, Anca Hermenean, Taras K. Oleksyk

## Abstract

**Background:** Population genomic projects are essential in the current drive to map the genome diversity of human populations across the globe. Various barriers persist hindering these efforts, and the lack of bioinformatic expertise and reproducible standardized population-scale analysis is one of the major challenges limiting their discovery potential. Scalable, automated, user-friendly pipelines can help researchers with minimum programming skills to tackle these issues without extensive training.

**Results:** *PopGenPlayground* (PGP), is a streamlined, single-command computation pipeline designed for human population genomics analysis based on *Snakemake* workflow management system. Developed to automate secondary analysis of a previously published national genome project, it leverages the publicly available genomic databases for comparative analysis and annotation of variant calls.

**Conclusions:** PGP presents a multi-platform robust population analysis pipeline, that reduces the time and the expertise levels to perform the main core of population analysis for a national genome project. PGP provides a comprehensive secondary analysis tool and can be used to perform analysis on a personal computer or using a remote high-performance computing platform.

## Background

The recent rise of population genomics can largely be attributed to major developments in sequencing technology and data analysis, and the exponential increase in the general volume of the data generated in the field (McGuire et al., 2020). In its nature (i.e. analysis of large datasets from multitude of samples), it is closely tied to the evolution of bioinformatics approaches and proliferation of modern computation techniques (Bartlett et al., 2017). Using new bioinformatics software and algorithms, researchers can efficiently analyze torrents of raw sequence data, identify variants, predict their functional effects, and examine patterns of genetic variation within and across populations of interest. Human population genomics projects are the necessary step towards providing information for and developing approaches in personalized medicine and advanced medical care (Polimanti et al., 2014).

Bioinformatics approaches facilitated the integration of different types of data in population genomics studies and opened the door to answering multi-domain research questions. For example, it is now possible to integrate genomic data with environmental, phenotypic, and geographic (Van Assche et al., 2015) data to study how various factors influence genetic variation within and among populations. This integrative approach has greatly enriched our understanding of population genomics. However, the growing complexity of modern research combined with the relative novelty and lack of standardized approaches for analyses establishes complex barriers to research groups starting their population genomics projects. Bioinformatics has made it possible but not necessarily made it accessible.

A typical bioinformatic analysis for any research study involves a multitude of tools and intermediary data manipulation steps, introducing an ever-growing bar in terms of expertise, forcing bioinformatics expert so specialize to narrow niches(Bartlett et al., 2017). One of the solutions to this problem aims to combine multiple steps and repetitive procedures in such analysis into a workflow. Streamlining, standardizing, and scaling up bioinformatics workflows, along with widespread knowledge sharing, can greatly improve the field’s accessibility. These initiatives have the potential to democratize genomics research, allowing a broader spectrum of researchers to participate in genome wide analysis of and reap the benefits of the big data approach.

### Purpose of PopGenPlayground

*PopGenPlayground* (PGP) is designed to facilitate comprehensive population genomics analysis using Variant Call Files (VCF). PGP suite is hosted on GitHub (Wolfsberger, 2023) and integrates a wide array of population genomics approaches (Table 1) into a single, user-friendly workflow. This pipeline automates key processes in population genomics analysis, including data wrangling, visualization, format conversion, and phasing, which reduces manual work and increases efficiency. *PGP* uses the *Snakemake* workflow management system (Köster et al., 2021) to ensure consistent computational environments throughout each analysis stage, limiting the impact of external variables. The workflow is scalable, capable of supporting high-performance computing environments, making it adaptable for large datasets. *PGP* features a simple configuration and does not require the user to have an advanced bioinformatics expertise to run.

**Table 1.** Analyses incorporated in the *PopGenPlayground* pipeline.

| Software\Tool | Class | Purpose | Reference |
| --- | --- | --- | --- |
| *BCFtools* | Data wrangling, VCF analysis | Provides an ability to merge and intersect Variant File Call, outputs statistics for comparative studies | (Danecek et al., 2021) |
| Whole Genome Sequencing Variant Call Files IGSR database | Comparative analysis | Provides populations to run the comparative analysis with other instruments | (Fairley et al., 2020) |
| *PLINK2* | Pedigree, Identity by descent/state, data wrangling, PCA test | Estimating inbreeding and relatedness, estimating structure, LD pruning, data conversion | (Chang et al., 2015) |
| *ADMIXTURE* | Clustering and characterizing admixture | Grouping individuals in clusters maximizing HW equilibrium and LD between loci | (Alexander et al., 2009) |
| *detectRUNS* (R package) | Runs of Homozygosity, Inbreeding based on runs of homozygosity | Estimating inbreeding and relatedness, detection of signals of selection | (Chang et al., 2015; Marras et al., 2015) |
| *Shapeit4* | Variant call phasing, | haplotype detection, missing data imputation | (Delaneau et al., 2008) |
| *Ensembl* VEP | Variant annotation | Combines multiple annotation approaches to assess the impact of variation on phenotype | (McLaren et al., 2016) |
| *Python* programming language and *Pandas* data analysis library | Data analysis tools | Report generation, data visualization, data wrangling | - |

### Implementation

The pipeline is developed with *Snakemake* workflow management system (Köster et al., 2021) using the Python-based rule definition language. The pipeline integrates various bioinformatics instruments, described in detail in **Table 1** to perform the analysis, and handles all the data wrangling and transformation steps internally using Unix shell commands, and Python programming language with *Pandas*, and associated data analysis libraries and the Fisher Exact Test (FET) statistics calculation module. The pipeline uses publicly available genomic databases. The International Genome Sample Resource or IGSR (Fairley et al., 2020) project was used for developing and evaluating the comparative analysis steps of the pipeline, databases associated with *Ensembl VEP* (McLaren et al., 2016) were used for genome annotations, while *NCBI ClinVar* data (Landrum et al., 2016) was integrated for annotation of clinical medical variation within the data.

Steps of the PGP analysis are defined as rules and direct the system on how to produce the desired output files. Dependencies of the rules are determined automatically, based on the input and output files defined. The integration of *Conda* package management system (“Anaconda Software Distribution,” 2020) handles the software dependencies of each workflow step. The pipeline adopts the “lazy” approach to fulfilling the rules, resolving them backwards from the final desired output file, and scanning the data folder for results of previous completed steps. This means an interruption of analysis can be resumed without the repetition of previously successful steps. Self-contained reports provide full transparency from results down to used steps, parameters, code, and software.

The pipeline uses minimalistic requirements in its configuration. After the initial installation of *Snakemake* and it’s requirements (Köster et al., 2021), a simple markup language format file is filled by the user. The input includes the VCF file of the population in study, and the three letter codes of populations from IGSR (Fairley et al., 2020) project. If there is no need to perfom all of the steps of the latter analysis, the configuration file can be given a simple binarry variable to enable-disable specific analytical steps. The pipeline follows the order of commands, provided in **Figure 1**.

**Figure 1.**
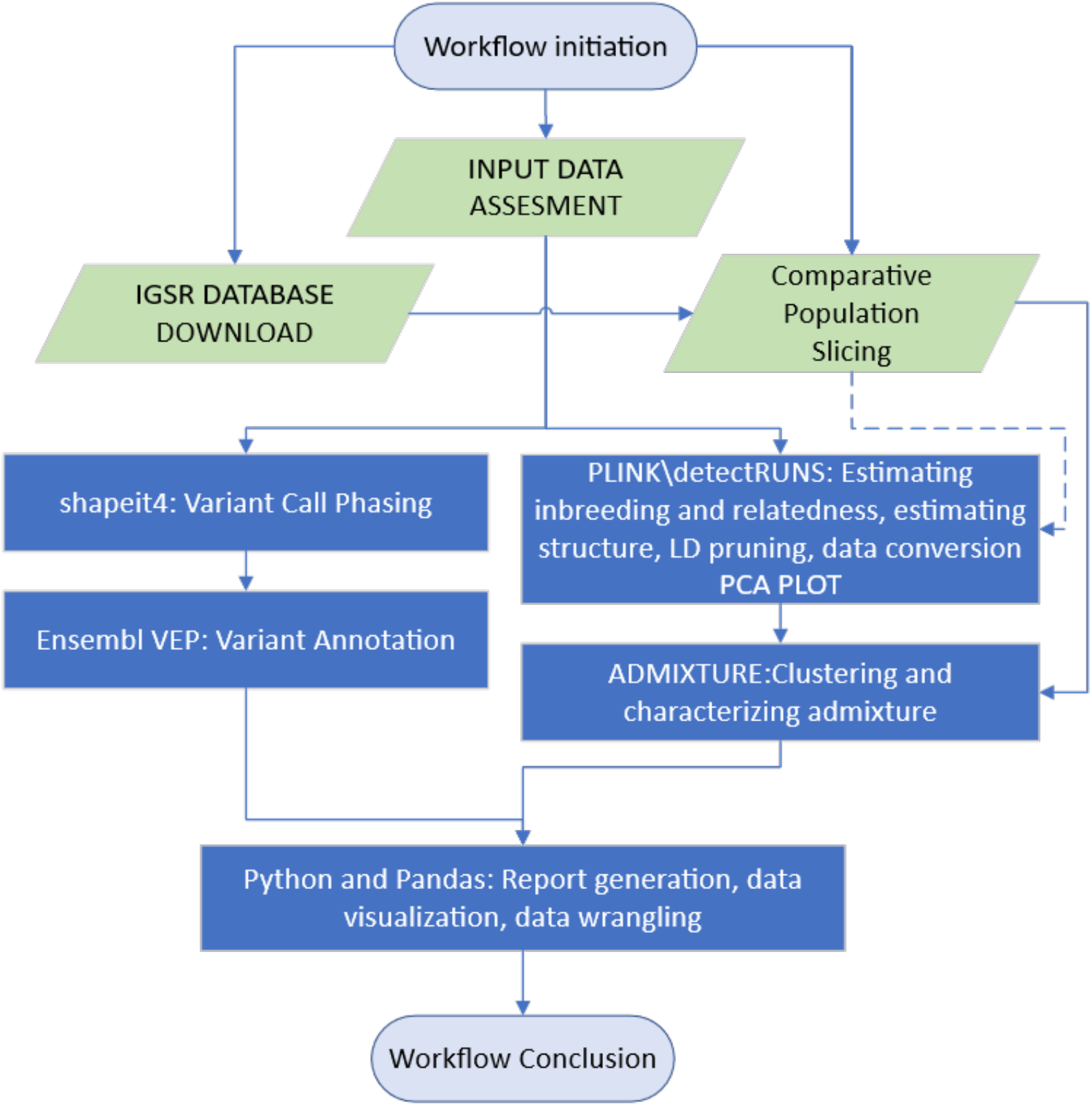
The schematic outline of the PopGenPlayground (PGP) pipeline.

### Results

Originally intended to replicate the results of the existing national genomics project analysis, the PGP pipeline can replicate the complete of the population genomics analysis performed in that study (Oleksyk et al., 2021). Using the pipeline replicates requires a minimal expertise, only the basic understanding of Unix shell-like command line. PGP leverages existing datasets and produces output providing valuable discoveries for the projects.

PGP provides a summary of variation population whole-genome sequences, offering insights into the total sequence reads, mean coverage, and variations such as SNPs, bi-allelic, multi-allelic, small indels, deletions, insertions, and structural variants. Additional table describes the summary annotation of different genomic elements, detailing the number of alleles and their distribution across various genomic locations like exons, introns, and intergenic regions. Lastly, it provides a dataset of comparative pairwise analysis of the population study, and all populations extracted from the IGSR (Fairley et al., 2020), highlighting the variants statistical different in their frequencies according to a Fisher Exact test.

Additionally, PGP produces various figures and associated datasets, using Python programming language, and various data analysis libraries. Clustering analysis results are summarized in PCA (**Figure 2.A**) and ADMIXTURE plots (**Figure 2.C**), illustrating the relationship between the samples within the population, and their comparative analysis with populations extracted from the IGSR. Variant calls with annotation and pairwise Fisher Exact Test (FET) information for allele frequencies are generated in a text table format. An example of a pre-publication figure output produced by PGP is shown in **Figure 2**.

**Figure 2.**
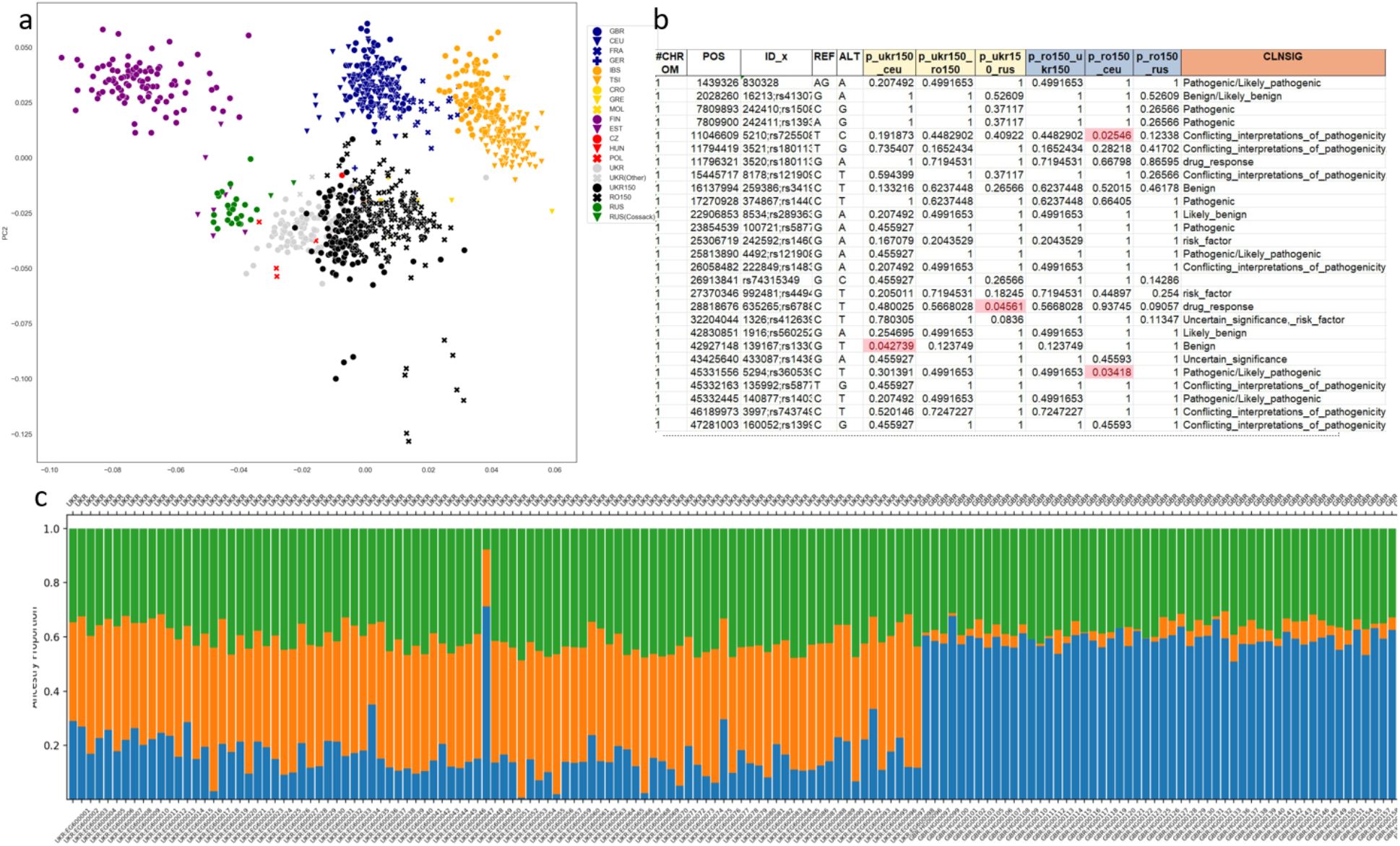
An example of the output produced by the *PopGenPlayground (PGP)*. a) PCA plot of populations in study combined with populations from the IGSR database (Fairley et al., 2020); b) Variant calls with annotation and pairwise Fisher Exact Test information for allele frequencies; c) A fragment of the Admixture plot of the population in study and IGSR-extracted population.

### Conclusions and future directions

To answer the rising demand in expertise and low accessibility of modern population genomics analysis approaches, we developed a bioinformatic pipeline *PopGenPlayground* (PGP). *PGP* is a robust bioinformatics tool providing user-friendly and efficient access to population genomics analyses. While utilizing the *Snakemake* workflow management system, PGP effectively combines diverse data types and integrates multiple bioinformatics tools, streamlining the analysis process. The design of the pipeline is based on previous expertise of analysis in a published national genomics project (Oleksyk et al., 2021) and integrates various public genomic databases for comparative analysis and variant annotation. Hosted on GitHub (Wolfsberger, 2023), PGP promotes collaboration and future growth of the pipeline. The field of population genomics is constantly changing, with new tools and important datasets being incorporated in them. Availability of new tools that can be incorporated into the analysis ensures the continuous development for the future growth of PGP pipeline, integrating more recent datasets, and expanding its analysis profile to include new instruments and data analyses.

## Declarations

### Authors Contributions

W.W.W. developed the pipeline and wrote the first draft. W.W.W. and K.S. prepared and analyzed the data. All other authors contributed ideas to the development of the pipeline and reviewed the ppeline and the manuscript. T.K.O. contributed to the original ideas, writing, and final editing of the manuscript.

### Software Availability

The complete pipeline and instructions for PopGenPlayground (PGP) pipeline use are available from the GitHub (*https://github.com/wwolfsberger/OU_popgen_playground*)(Wolfsberger, 2023)

### List of Abbreviations

PGP: PopGenPlayground; VCF: Variant Call Format.

### Competing Interests

The authors declare that they have no competing interests.

### Funding

Funding for the project was provided by 2SOFT/1.2/48 project “*Partnership for Genomic Research in Ukraine and Romania*” by the *Joint Operational Programme Romania-Ukraine*, through the *European Neighbourhood Instrument* (ENI).

## Acknowledgements

This instrument is part of the developing infrastructure for bioinformatics in Ukraine. We thank all the participants of the *BioinformaticsForUkraine*.*com* and the *Genome Diversity in Ukraine Consortium* who worked with us on developing tools for this project.

## Notes

https://github.com/wwolfsberger/OU_popgen_playground

### Competing Interest Statement

The authors have declared no competing interest.

## References

Alexander, D. H., Novembre, J., & Lange, K. (2009). Fast model-based estimation of ancestry in unrelated individuals. Genome Research, 19(9), 1655–1664. 10.1101/GR.094052.109

Anaconda Software Distribution. (2020). In Anaconda Documentation. Anaconda Inc. https://docs.anaconda.com/

Bartlett, A., Penders, B., & Lewis, J. (2017). Bioinformatics: Indispensable, yet hidden in plain sight? BMC Bioinformatics, 18(1), 1–4. 10.1186/S12859-017-1730-9/METRICS

Chang, C. C., Chow, C. C., Tellier, L. C. A. M., Vattikuti, S., Purcell, S. M., & Lee, J. J. (2015). Second-generation PLINK: Rising to the challenge of larger and richer datasets. GigaScience, 4(1), 7. 10.1186/s13742-015-0047-8

Danecek, P., Bonfield, J. K., Liddle, J., Marshall, J., Ohan, V., Pollard, M. O., Whitwham, A., Keane, T., McCarthy, S. A., & Davies, R. M. (2021). Twelve years of SAMtools and BCFtools. GigaScience, 10(2). 10.1093/GIGASCIENCE/GIAB008

Delaneau, O., Coulonges, C., & Zagury, J. F. (2008). Shape-IT: New rapid and accurate algorithm for haplotype inference. BMC Bioinform., 9, 1–14. 10.1186/1471-2105-9-540

Fairley, S., Lowy-Gallego, E., Perry, E., & Flicek, P. (2020). The International Genome Sample Resource (IGSR) collection of open human genomic variation resources. Nucleic Acids Research, 48(D1), D941–D947. 10.1093/NAR/GKZ836

Köster, J., Mölder, F., Jablonski, K. P., Letcher, B., Hall, M. B., Tomkins-Tinch, C. H., Sochat, V., Forster, J., Lee, S., Twardziok, S. O., Kanitz, A., Wilm, A., Holtgrewe, M., Rahmann, S., & Nahnsen, S. (2021). Sustainable data analysis with Snakemake. F1000Research 2021 10:33, 10, 33. 10.12688/f1000research.29032.2

Landrum, M. J., Lee, J. M., Benson, M., Brown, G., Chao, C., Chitipiralla, S., Gu, B., Hart, J., Ho?man, D., Hoover, J., Jang, W., Katz, K., Ovetsky, M., Riley, G., Sethi, A., Tully, R., Villamarin-Salomon, R., Rubinstein, W., & Maglott, D. R. (2016). ClinVar: public archive of interpretations of clinically relevant variants. Nucleic Acids Research, 44(D1), D862–D868. 10.1093/nar/gkv1222

Marras, G., Gaspa, G., Sorbolini, S., Dimauro, C., Ajmone-Marsan, P., Valentini, A., Williams, J. L., & MacCiotta, N. P. P. (2015). Analysis of runs of homozygosity and their relationship with inbreeding in ?ve cattle breeds farmed in Italy. Animal Genetics, 46(2), 110–121. 10.1111/AGE.12259

McGuire, A. L., Gabriel, S., Tishkoff, S. A., Wonkam, A., Chakravarti, A., Furlong, E. E. M., Treutlein, B., Meissner, A., Chang, H. Y., López-Bigas, N., Segal, E., & Kim, J.-S. (2020). The road ahead in genetics and genomics. Nature Reviews Genetics, 21(10), 581–596. 10.1038/s41576-020-0272-6

McLaren, W., Gil, L., Hunt, S. E., Riat, H. S., Ritchie, G. R. S., Thormann, A., Flicek, P., & Cunningham, F. (2016). The Ensembl Variant Effect Predictor. Genome Biology. 10.1186/s13059-016-0974-4

Oleksyk, T. K., Wolfsberger, W. W., Weber, A. M., Shchubelka, K., Oleksyk, O. T., Levchuk, O., Patrus, A., Lazar, N., Castro-Marquez, S. O., Hasynets, Y., Boldyzhar, P., Neymet, M., Urbanovych, A., Stakhovska, V., Malyar, K., Chervyakova, S., Podoroha, O., Kovalchuk, N., Rodriguez-Flores, J. L., … Smolanka, V. (2021). Genome diversity in Ukraine. GigaScience, 10(1), 1–14. 10.1093/GIGASCIENCE/GIAA159

Van Assche, R., Broeckx, V., Boonen, K., Maes, E., De Haes, W., Schoofs, L., & Temmerman, L. (2015). Integrating -Omics: Systems Biology as Explored Through C. elegans Research. Journal of Molecular Biology, 427(21), 3441–3451. 10.1016/J.JMB.2015.03.015

Wolfsberger, W. W. (2023). PopGenPlayground (0.1). https://github.com/wwolfsberger/OU_popgen_playground.

